# Utilization efficiency of human milk oligosaccharides by human-associated *Akkermansia* is strain-dependent

**DOI:** 10.1101/2021.07.26.453919

**Authors:** Estefani Luna, Shanthi G. Parkar, Nina Kirmiz, Stephanie Hartel, Erik Hearn, Marziiah Hossine, Arinnae Kurdian, Claudia Mendoza, Katherine Orr, Loren Padilla, Katherine Ramirez, Priscilla Salcedo, Erik Serrano, Biswa Choudhury, Mousumi Paulchakrabarti, Craig T. Parker, Steven Huynh, Kerry Cooper, Gilberto E. Flores

## Abstract

*Akkermansia muciniphila* are mucin degrading bacteria found in the human gut and are often associated with positive human health. However, despite being detected as early as one month of age, little is known about the role of *Akkermansia* in the infant gut. Human milk oligosaccharides (HMOs) are abundant components of human milk and are structurally similar to the oligosaccharides that comprise mucin, the preferred growth substrate of human-associated *Akkermansia*. A limited subset of intestinal bacteria has been shown to grow well on HMOs and mucin. We therefore examined the ability of genomically diverse strains of *Akkermansia* to grow on HMOs. First, we screened 85 genomes representing the four known *Akkermansia* phylogroups to examine their metabolic potential to degrade HMOs. Furthermore, we examined the ability of representative isolates to grow on individual HMOs in a mucin background and analyzed the resulting metabolites. All *Akkermansia* genomes were equipped with an array of glycoside hydrolases associated with HMO-deconstruction. Representative strains were all able to grow on HMOs with varying efficiency and growth yield. Strain CSUN-19 belonging to the AmIV phylogroup, grew to the highest level in the presence of fucosylated and sialylated HMOs. This activity may be partially related to the increased copy numbers and/or the enzyme activities of the α-fucosidases, α-sialidases, and β-galactosidases. Utilization of HMOs by strains of *Akkermansia* suggests that ingestion of HMOs by an infant may enrich for these potentially beneficial bacteria. Further studies are required to realize this opportunity and deliver long-lasting metabolic benefits to the human host.

**Importance:** Human milk oligosaccharides (HMOs) are utilized by a limited subset of bacteria in the infant gut. *Akkermansia* are detected in infants as young as one month of age and are thought to contribute to the HMO deconstruction capacity of the infant. Here, using phylogenomics, we examined the genomic capacity of different *Akkermansia* phylogroups to potentially deconstruct HMOs. Furthermore, we experimentally showed that strains from all the currently known phylogroups of *Akkermansia* can deconstruct all the major types of HMOs, albeit with different utilization efficiencies. This study thus examines *Akkermansia*-HMO interactions that can potentially influence the gut microbial ecology during the first 1,000 days of life - a critical phase for the development of the gut microbiome and infant health.

This study will be of interest to a wide range of scientists from microbiologists, glycochemists/glycobiologists, to functional food developers investigating *Akkermansia* as probiotics or functional foods containing milk oligosaccharides as prebiotics.

## Introduction

*Akkermansia muciniphila* is a mucin-degrading specialist that colonizes the mucus layer of the human gastrointestinal tract.^1^ Paradoxically, *Akkermansia* also promote mucus production by enhancing the differentiation of gut epithelial cells, thereby influencing mucosal homeostasis.^2^ Numerous positive associations have been observed between this bacterial lineage and human health. In adults, a decreased abundance of *Akkermansia* is associated with metabolic impairments,^3^ ulcerative colitis,^4^ and inflammatory bowel disease.^5^ In infants, a decrease in mucosal residents such as *Akkermansia* is associated with a compromised immune system and the development of atopic dermatitis.^6^

The mechanisms by which *A. muciniphila* benefit human health appears to be directly linked to its ecological niche along the human gastrointestinal tract. Specifically, *A. muciniphila* colonizes the oxic-anoxic interface of the mucus layers adjacent to host epithelial cells where they degrade host-produced mucins.^7^ Mucins are the main structural components of mucus and are composed of polypeptide chains rich in serine, threonine, and proline residues that are *O*-linked to a variety oligosaccharides.^8^ These oligosaccharide side chains are comprised of N-acetylgalactosamine (GalNAc), N-acetylglucosamine (GlcNAc), galactose, and are capped with N-acetylneuraminic acid (Neu5Ac; sialic acid), fucose, or sulfate. *Akkermansia* can utilize mucins as their sole carbon and nitrogen source, generating metabolites such as acetate and succinate, and propionate in the presence of vitamin B12.^9,10^ Co-occurring members of the gut microbiome convert some of the acetate produced to butyrate.^11^ Together, these organic acids fuel colonocytes and act as signaling molecules helping to maintain an overall anti-inflammatory tone in the gut.^12^ In addition to producing ant-inflammatory metabolites, *A. muciniphila* produces an extracellular surface protein, coded by Amuc_1100, that interacts directly with Toll-Like Receptors on host epithelial cells.^13,14^ This interaction results in the production of specific anti-inflammatory cytokines including IL-10 that leads to an improvement in overall gut barrier function.^13^

Building upon previous work by Guo and colleagues,^15^ we recently performed a comparative genomic analysis of 75 *Akkermansia* genomes to define the genomic and functional landscape of this lineage. This analysis identified at least four distinct phylogroups AmI-AmIV, with the type strain *A. muciniphila* Muc^T^, belonging to the AmI phylogroup. Additionally, this work showed that the *Akkermansia* phylogroups had differing functional potentials including *de novo* biosynthesis of vitamin B12 by members of the AmII and AmIII phylogroups.^10^

Continuing to explore the genomic and metabolic diversity of human-associated *Akkermansia*, we next wanted to determine if host-produced glycans, other than those in mucin, could support growth of the various *Akkermansia* phylogroups. Because of the compositional and structural similarities between the oligosaccharides found in mucin and human milk, we focused on human milk oligosaccharides (HMOs).^8,16,17^ Human milk contains 5-15 g/L HMOs, of which 50-80% are fucosylated, and 10-20% are sialylated.^16^ Although HMOs are present in milk as a pool of over 200 diverse structures, they are composed of only five monosaccharides: glucose, galactose, fucose, N-acetylglucosamine (GlcNAc), and sialic acid.^16^ These oligosaccharides contain a lactose core at the reducing end that is extended with building block monosaccharides via glycosidic linkages. In human milk, fucose can be attached via α1-2, α1-3, and α1-4 linkages, and sialic acid can be attached via α2-3 and α2-6 linkages. Simple, abundant, and routinely studied HMO structures include lacto-N-tetraose (LNT), lacto-N-neotetraose (LNnT), 2’-fucosyllactose (2’-FL), 3-fucosyllactose (3-FL), 6’-sialyllactose (6’-SL), and 3’-sialyllactose (3’-SL).^18^

The oligosaccharides found in human milk are not digestible by the developing infant and reach the intestine intact.^19^ Once there, HMOs have a variety of functions including providing protection from pathogens, playing a role in modulation of gut epithelial cells, and enriching for a beneficial microbiota.^20–22^ Several studies have screened HMO consumption by various intestinal commensals and have identified a limited group of bacteria, primarily *Bifidobacterium* and select *Bacteroides*, with this ability.^23–25^ One *Akkermansia* strain, belonging to phylogroup AmI, (i.e., *A. muciniphila* Muc^T^) has recently been shown to grow on human milk and select HMOs using a repertoire of glycoside hydrolase (GH) enzymes.^26^ In this current study, we expand our understanding of this HMO-degrading capacity of human-associated *Akkermansia* beyond the one phylogroup. We hypothesized that *Akkermansia* from the different phylogroups will differ in their ability to metabolize HMOs, and these differences are related to their genomic composition. To investigate the ability of *Akkermansia* to grow on select HMO, we first took a comparative genomics approach focusing on the presence and abundance of genes coding for glycoside hydrolase enzymes known to be involved in HMO catabolism. We then performed comparative growth experiments and demonstrated robust growth of representative strains from each phylogroup in a basal medium supplemented with five individual pure HMOs, in a background of mucin – thus simulating the carbon sources available in the infant gut environment. These findings expand the known metabolic capabilities of human-associated *Akkermansia* and point to further functional differences among the genomically distinct phylogroups.

## Results

### Isolation, identification, and genomics

In total, 17 human-associated *Akkermansia* were isolated from healthy adults, 10 from males, and 7 from females (Supplemental Table S2). Phylogenetic analyses of nearly complete 16S rRNA gene sequences from each isolate revealed three well-supported clades with the AmIII phylogroup nested within the AmII phylogroup (Figure 1A). At least one isolate was obtained from the four known human-associated phylogroups.^10^ Ten of the 17 isolates treed within the AmI phylogroup, followed by four AmII, two AmIV, and one AmIII.

**Figure 1.**
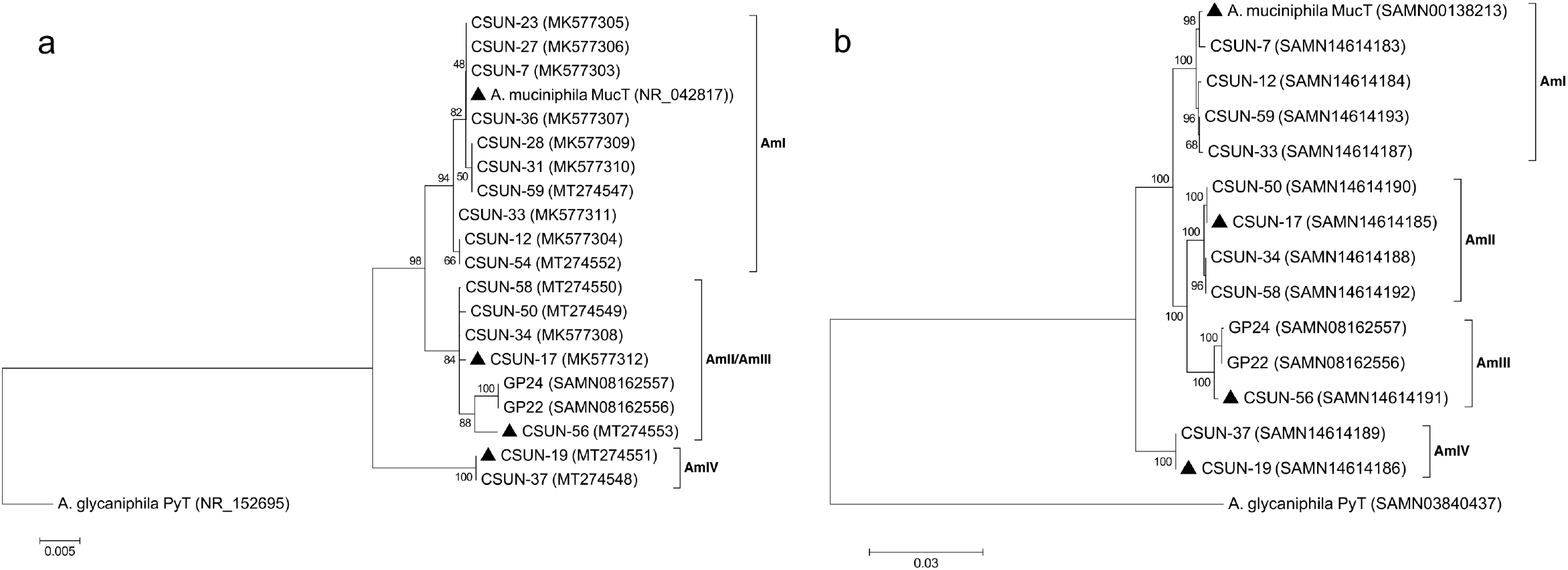
Phylogenetic relationship of *Akkermansia* isolates based on near-full length 16S rRNA gene sequences (**A**) and concatenation of 49 ribosomal protein coding genes obtained from draft genomes (**B**). Both trees are rooted using the only other named species of the genera, *A. glycaniphila* Py^T^. Isolates with triangles were used in HMO growth experiments. GP22 and GP24 in the AmIII phylogroup are from Guo and colleagues ^15^ and are included because only one AmIII representative is available in our culture collection. Both trees were generated in MEGA7 ^67^ using the maximum likelihood method and numbers at nodes indicate bootstrap values for 100 replicates. The tree in **A** was generated considering only unambiguously aligned nucleotide positions (n=1,305). For **B**, a total of 7,327 amino acid positions across 49 protein-coding genes were used. Both trees are drawn to scale with branch lengths measured in the number of substitutions per site.

Using the 16S rRNA tree as a guide, we selected 11 of the isolates spanning each phylogroup for genomic sequencing. Characteristics of these draft genomes are presented in Table 1. Of the new isolates, draft genome size ranged from 2.86 Mb (CSUN-56, AmIII) to 3.15 Mb (CSUN-19, AmIV) with 2,658 to 3,111 coding sequences (CDs), respectively, as compared with 2.67 Mb genome size and 2,576 CDs in *A. muciniphila* Muc^T^. Across phylogroups, approximately 52% of CDs could be assigned a function, on average. Resolution of the AmIII phylogroup was improved with phylogenomic analysis that included 49 protein-coding genes (Figure 1B).

**Table 1.** Genomic properties of 11 human-associated *Akkermansia* isolates. For comparison, the fasta sequence of the type strain *Akkermansia muciniphila* Muc^T^ was downloaded from GenBank accession number CP001071.1 and analyzed identically to the new isolates.

| Strain | Contigs | GC Content (%) | Genome Length (bp) | CDs | tRNA | rRNA | Hypothetical proteins | Proteins with assignments | EC number assignments | GO assignments | KEGG pathways |
| --- | --- | --- | --- | --- | --- | --- | --- | --- | --- | --- | --- |
| *A. muciniphila* Muc<sup>T</sup> (AmI) | 1 | 55.8 | 2,664,102 | 2,576 | 53 | 3 | 1,072 | 1,504 | 620 | 529 | 459 |
| *Akkermansia* CSUN-7 (AmI) | 52 | 55.1 | 2,875,736 | 2,880 | 50 | 3 | 1,377 | 1,503 | 623 | 530 | 464 |
| *Akkermansia* CSUN-12 (AmI) | 56 | 55.3 | 2,810,203 | 2,823 | 50 | 3 | 1,307 | 1,516 | 632 | 538 | 465 |
| *Akkermansia* CSUN-33 (AmI) | 49 | 55.3 | 2,833,117 | 2,853 | 50 | 3 | 1,307 | 1,546 | 633 | 541 | 469 |
| *Akkermansia* CSUN-59 (AmI) | 65 | 55.2 | 2,942,175 | 3,010 | 50 | 3 | 1,472 | 1,538 | 628 | 535 | 466 |
| *Akkermansia* CSUN-17 (AmII) | 29 | 58.2 | 2,999,178 | 2,856 | 49 | 3 | 1,354 | 1,502 | 641 | 542 | 474 |
| *Akkermansia* CSUN-34 (AmII) | 87 | 57.8 | 3,024,116 | 2,949 | 47 | 3 | 1,451 | 1,498 | 640 | 538 | 473 |
| *Akkermansia* CSUN-50 (AmII) | 23 | 58.2 | 3,005,559 | 2,842 | 49 | 3 | 1,365 | 1,477 | 632 | 533 | 470 |
| *Akkermansia* CSUN-58 (AmII) | 71 | 57.8 | 3,087,515 | 2,988 | 49 | 3 | 1,489 | 1,499 | 637 | 535 | 472 |
| *Akkermansia* CSUN-56 (AmIII) | 48 | 58.5 | 2,860,685 | 2,658 | 48 | 3 | 1,246 | 1,412 | 612 | 518 | 462 |
| *Akkermansia* CSUN-19 (AmIV) | 89 | 56.6 | 3,149,202 | 3,111 | 49 | 3 | 1,656 | 1,455 | 628 | 535 | 472 |
| *Akkermansia* CSUN-37 (AmIV) | 72 | 56.7 | 3,142,630 | 3,077 | 49 | 3 | 1,631 | 1,446 | 624 | 531 | 469 |

CDs = coding sequences, EC = Enzyme Classification, GO = Gene Ontology, KEGG = Kyoto Encyclopedia of Genes and Genomes

To investigate the carbohydrate degrading potential of the *Akkermansia* strains, 85 genomes, including the 11 isolates from this study, were annotated against the CAZy database,^27^ using dbCAN.^28,29^ We first took a global look at all annotated GH families and found significantly less GH annotations in genomes from the AmI phylogroup compared to other phylogroups (Kruskal-Wallis, χ^2^ = 55.128, P < 0.0001, Supplemental Figure S2). Further, we identified consistent similarities and differences in the complement of GH annotations within and between each phylogroup (Figure 2). With a few minor exceptions, these similarities and differences in GH counts resulted in the clustering of genomes into their respective phylogroups as evidenced by the dendrogram along the y-axis in Figure 2.

**Figure 2.**
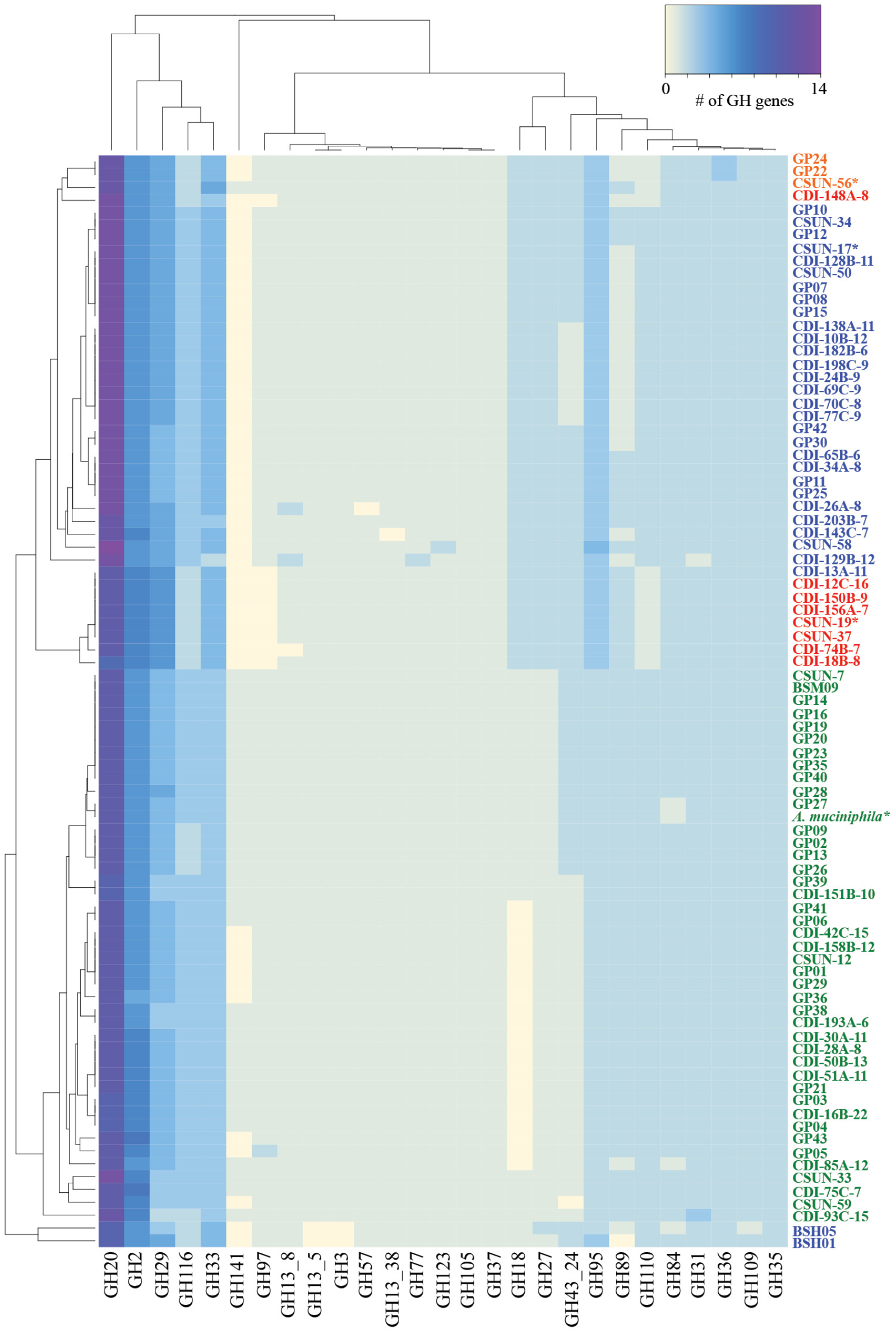
Human-associated *Akkermansia* possess different complements of glycoside hydrolase (GH) genes potentially impacting their carbohydrate degrading capabilities. The heat map shows the counts of different GH families present in the draft genome of 85 total *Akkermansia* genomes. Genomes labeled with ‘CSUN’ prefixes are isolates from this work while the ‘CDI’ genomes are from metagenome assembled genomes ^10^ and the ‘GP’ or ‘BSM’ genomes are from isolates from Guo and colleagues ^15^. Each genome is colored by phylogroup affiliation with green = AmI, blue = AmII, orange = AmIII, and red = AmIV. Only three genomes (CDI-148A-8, BSH05, and BSH01) tree outside of their phylogroup affiliation based on the GH content. Genomes with asterisks were used in the HMO growth experiments.

Next, since we were interested in the ability of *Akkermansia* to degrade HMO, we focused on HMO-associated GH families previously identified in other organisms.^26,30–33^ With this approach, we identified differences in the copy number of several GH families that are associated with degradation of HMO-glycans; α-fucosidases, α-sialidases, β-galactosidases, N-acetyl β-hexosaminidases (Table 2). Most of these genes were also found to possess a signal peptide (Supplementary Excel Data 1), which is indicative of encoding for extracellular enzymes.^34,35^ Of note was the high number of GH20 genes as compared with any other GH gene in all the genomes. The number of putative α-fucosidases (GH29, GH95 and GH141) and N-acetyl β-hexosaminidases (GH18, GH20, GH84 and GH109) also varied across phylogroups (lowest for AmI including the strain tested here, *A. muciniphila* Muc^T^). Of the four strains investigated for HMO catabolic capacity in this study, the CSUN-19 (AmIV phylogroup) and CSUN-56 (AmIII) strains showed 9 fucosidase annotations as compared with 8 for CSUN-17 (AmII) and 7 for *A. muciniphila* Muc^T^ (AmI; Table 2).

**Table 2.** Copy number of several human milk oligosaccharide-associated glycoside hydrolase (GH) families in representative strains from the different *Akkermansia* phylogroups. The average copy number for each phylogroup is given in parentheses.

| Glycoside |  | *A. muciniphila* | CSUN-17 | CSUN-56 | CSUN-19 |
| --- | --- | --- | --- | --- | --- |
| Hydrolase | Enzyme activity | Muc<sup>T</sup> (AmI) | (AmII) | (AmIII) | (AmIV) |
| Family |  |  |  |  |  |
| GH2 | β-galactosidase (or similar) | 6 (6.3) | 6 (6) | 6 (6) | 7 (6.9) |
| GH16 | β-galactosidase (or similar) | 3 (2.9) | 3 (2.9) | 2 (2) | 2 (2) |
| GH18 | Chitinase; endo-β-N-acetylglucosaminidase (or similar) | 1 (0.5) | 2 (1.9) | 2 (2) | 2 (2) |
| GH20 | β-hexosaminidase; lacto-N-biosidase; β-N-acetylglucosaminidase | 11 (11) | 13 (12.8) | 12 (12) | 11 (11) |
| GH29 | α-L-fucosidase | 4 (3.8) | 5 (4.7) | 5 (5) | 6 (5.9) |
| GH33 | Sialidase (or similar) | 3 (3) | 4 (3.9) | 5 (4.3) | 4 (3.9) |
| GH35 | β-galactosidase; exo-β-glucosaminidase | 2 (2) | 2 (2) | 2 (2) | 2 (2) |
| GH84 | N-acetyl β-glucosaminidase; hyaluronidase | 1 (1.9) | 2 (2) | 2 (2) | 2 (2) |
| GH95 | α-L-fucosidase; α-L-galactosidase | 2 (2) | 3 (3) | 3 (3) | 3 (3) |
| GH109 | α-N-acetylgalactosaminidase; β-N-acetylhexosaminidase | 2 (2) | 2 (2) | 2 (2) | 2 (2) |
| GH141 | α-L-fucosidase; xylanase | 1 (0.8) | 0 (0) | 1 (0.3) | 0 (0) |

### Utilization of HMOs

Representatives of each phylogroup were tested for their ability to grow on HMO in the presence of mucin. After 48 h of incubation, all strains tested grew to higher ODs in the HMO (or lactose)-supplemented mucin medium compared to growth in medium lacking HMOs (Figure 3). Growth yield varied across strains on media with 2’-FL, 3-FL, LNnT, and 6’-SL, but not LNT or lactose (P < 0.05, ANOVA). Post-hoc comparisons revealed that strain CSUN-19, representing the AmIV phylogroup, showed the greatest growth in comparison to the other strains, with significant increases compared with *A. muciniphila* Muc^T^ in 2’-FL, 3-FL, and 6’-SL; and with CSUN-56 in 2’-FL, 3-FL, and LNnT (Figure 3).

**Figure 3.**
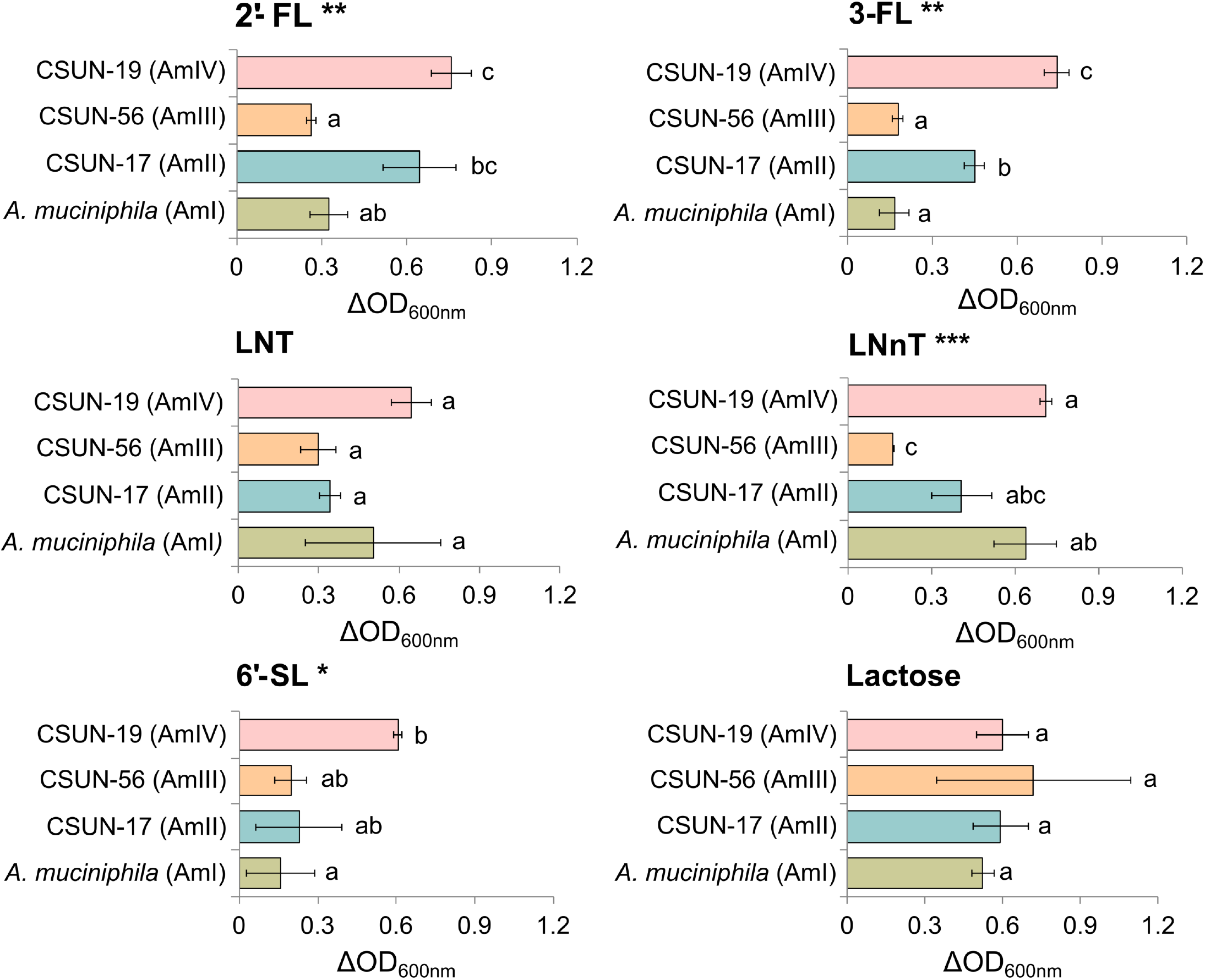
Representative strains from the four *Akkermansia* phylogroups were incubated in a mucin-containing medium alone or supplemented with 20 mM of individual human milk oligosaccharides or lactose. The experiment was conducted in triplicate and repeated at least two times. The difference in OD_600nm_ from the growth in the mucin-containing medium alone was used to plot the bacterial growth for each strain. Values are expressed as average +/- standard deviation. The ANOVA tests reveal significant effects denoted by **, P < 0.01 and * P < 0.05 with the substrates 2’-fucosyllactose (2’-FL), 3-fucosyllactose (3-FL), lacto-N-neotetraose (LNnT), and 6’-sialyllactose (6’-SL) but not with lacto-N-tetraose (LNT) and lactose. Pairwise comparisons within each substrate using Tukey’s Honestly Significant Difference test reveal significant differences between the phylogroups (P < 0.05) - means showing letters in common are not significantly different.

To confirm HMO utilization, we measured the concentrations of HMOs (2’-FL, LNT and 6’-SL) and their sugar constituents (except GlcNAc for LNT) before and after 48 h of incubation (Figure 4a and 4b). In addition to the difference in growth yield, the difference in the % HMO utilized also varied across strains (P < 0.05). For 2’-FL, strains representing the AmI, AmII, and AmIII phylogroups utilized greater than 93% of the available HMO, while CSUN-19 (AmIV) utilized just over 64% despite having the highest growth yield as measured by the change in OD_600nm_. Nearly all of the fucose liberated from 2’-FL was removed from the medium within 48 h by all the strains, while the lactose backbone accumulated in the culture medium of all strains except CSUN-19 (AmIV) (Figure 4c). Degradation of LNT ranged from 25.4 −78.6% across tested strains with CSUN-17 (AmII) utilizing the least and *A. muciniphila* Muc^T^ (AmI) utilizing the most. In contrast to growth on 2’-FL, most of the lactose from LNT was consumed across strains (Figure 4d). Similar to LNT, there was a wide range of 6’-SL utilization across strains (P < 0.001), ranging from 29.3% (CSUN-17, AmII) to 89.2% (CSUN-19, AmIV). In the case of 6’-SL, CSUN-19 showed the greatest growth, while *A. muciniphila* Muc^T^ showed the least growth, and yet the % substrate utilized (80%) showed no significant difference and was significantly higher than the ~50% and ~30% utilization seen with CSUN-56 and CSUN-17, respectively. In all strains, sialic acid accumulated in the culture media and was not consumed when liberated from 6’-SL (Figure 4e).

**Figure 4.**
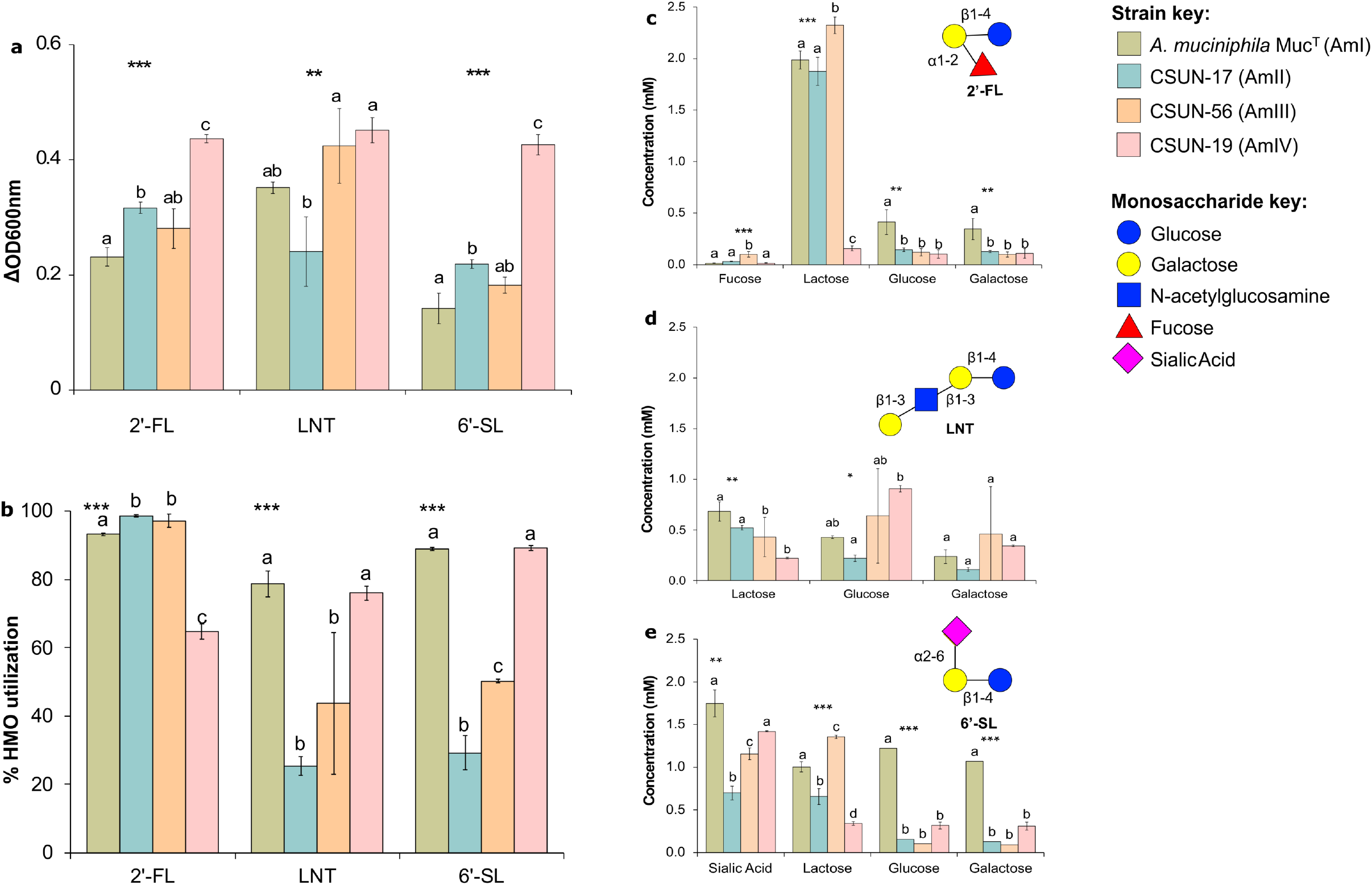
Representative strains from the four *Akkermansia* phylogroups were incubated in a mucin-containing medium alone or supplemented with 4 mM of individual human milk oligosaccharides (HMOs) or lactose. The experiment was conducted in triplicate and repeated three times. (a) The difference in growth in the HMO-supplemented medium from the growth in the mucin-containing medium alone was used to plot the bacterial growth for each strain. (b) The concentrations of the original substrate analyzed were used to calculate the percentage of the HMO utilized. (c,d,e) The concentrations of the metabolites obtained after the deconstruction of the 2’-fucosyllactose (2’-FL), lacto-N-tetraose (LNT) and 6’-sialyllactose (6’-SL) respectively are expressed as average +/- standard deviation. Statistical analysis revealed significant effects between substrates (a,b) and strains (b,c,d), denoted by ***, P < 0.001; **, P < 0.01 and *, P < 0.05. Pairwise comparisons using Tukey’s Honestly Significant Difference test were also performed with P < 0.05, and means showing letters in common are not significantly different.

## Discussion

*Akkermansia* are largely considered beneficial members of the human gut microbiome and are currently of significant interest for their therapeutic potential.^36^ Until recently, however, all research involving these promising bacteria focused on a single species, *A. muciniphila* Muc^T^ belonging to the AmI phylogroup. Here, we continue to build upon recent work by ourselves and others describing genomic and functional diversity within this lineage.^10,37^ Specifically, we show genomically diverse strains possess different complements of GH genes that encode enzymes catalyzing the deconstruction of HMOs into constituent mono- and disaccharides. Furthermore, we demonstrate that four different *Akkermansia* strains representing the four known phylogroups can deconstruct HMOs, with this biological activity varying across strains. These differences in genomic and functional traits of the human-associated *Akkermansia*, along with diversity in the substrates that are presented to the gut bacteria in the form of breast milk or supplemented infant milk formula, potentially impact how and when *Akkermansia* colonize the human gastrointestinal tract. For example, the ability to utilize HMOs efficiently could provide a competitive advantage for the early colonization of the infant gut with human-associated *Akkermansia* in a strain-dependent manner. *Akkermansia* are key contributors to the infants’ glycan-metabolizing capacity as early as 4 months of age,^33^ and may therefore play a critical role in establishing a foundation of metabolic fitness in the naïve microbiome. Taken together, these findings expand the known metabolic niche and interaction network of *Akkermansia* in the human gut early in life.

Bacterial growth studies have demonstrated that relatively few gut bacteria grow well on HMOs, the exceptions being bifidobacteria and select *Bacteroides*, both dominant members of the infant gut.^24,25^ Both bifidobacteria and *Bacteroides* employ an array of glycoside hydrolases including fucosidases (GH29 and GH95), sialidases (GH33), galactosidases (GH2 and GH16), lacto-N-biosidases (GH20), and hexosaminidases (GH20) to deconstruct HMO linkages.^38–44^ Our phylogenomic characterization of the *Akkermansia* genomes shows that the various strains from the four *Akkermansia* phylogroups possess a wealth of these same gene annotations, albeit in differing abundances, that could be used for the deconstruction of either HMO or mucin. Bifidobacteria employ two major strategies to hydrolyze HMOs.^31,42^ Infant-associated *Bifidobacterium infantis*, *Bifidobacterium breve*, and *Bifidobacterium longum* primarily consume HMOs by employing intracellular glycoside hydrolases to deconstruct the HMO structures.^41,42,45–47^ Using an alternative strategy, *Bifidobacterium bifidum* extracellularly process the HMO via an array of membrane associated glycoside hydrolases. ^31,48^ *Bacteroides* spp. harbor polysaccharide utilization loci (PULs) that encode a diverse array of glycosidases capable of breaking down host-produced and plant-derived polysaccharides.^44,49^ *Bacteroides* are hypothesized to bind HMOs on the cell surface followed by hydrolysis of the HMOs and import of the resultant oligo-saccharides for further breakdown. They co-opt their mucin-utilization PULs to deconstruct and utilize HMOs with varying efficiency depending on the strain. *B. fragilis* are the most efficient preferring HMOs with a high degree of polymerization and non-fucosylated HMOs over fucosylated HMOs,^24^ and even utilize the sialic acid generated after deconstruction of sialylated HMOs.^44^ *Akkermansia* do not have the typical PUL genomic organization seen in the *Bacteroides*, but they do appear to harness extracellular GHs either in the periplasmic space or outside of the cell altogether to cleave monosaccharides or disaccharides from mucin or HMOs.^9,26^ In agreement, the majority of our GH annotations included signal peptide sequences indicative of export outside of the cytoplasmic membrane. Extracellular cleavage of HMO (and mucin) results in the liberation of monosaccharides and disaccharides that enables cross-feeding by other members of the gut microbiome.^11^ In the context of the infant gut, this cross-feeding could help facilitate colonization of new members to the gut community that are encountered as infants grow and consume new foods, aiding in the maturation of the gut microbiome in the early years of life.

In addition to the cross-feeding on sugars liberated from host substrates, members of the gut microbiome feed off fermentation waste products produced by *Akkermansia*. In the case of fucosylated substrates such as 2’-FL, a distinct metabolite of fucose fermentation is 1,2-propanediol.^26^ Several bacterial genera including both beneficial (*Lactobacillus* spp., *Eubacterium hallii*) and pathogenic (*Salmonella*) bacteria, can grow on 1,2-propanediol in a vitamin B12-dependent manner.^50,51^ Given our recent work showing that the AmII and AmIII phylogroups synthesize vitamin B12^10^, these findings indicate the possibility of *Akkermansia*-driven syntrophic interactions that are likely phylogroup-specific. This is particularly relevant as the gut microbiome of exclusively breast-fed infants has a decreased capacity for *de novo* synthesis of vitamin B12, compared with formula-fed infants.^52^ Therefore, if *Akkermansia* are to be used therapeutically, then it will be important to consider the strain to be used in the context of the host’s age and health status. Alternatively, if *Akkermansia* are already present in the host, it will be important to know which strain is present to better predict the outcome of any microbiome or dietary intervention.

Several studies have detected *Akkermansia* in stool of infants as early as one-month after birth, in most one-year olds,^53^ and even in human colostrum and milk,^54,55^ demonstrating that it colonizes the gut early in life and providing a possible route of inoculation. Two separate studies found direct associations between the abundance of *Akkermansia* and fucosylated HMO in human milk, suggesting that fucosylated HMO may help enrich for *Akkermansia* in the gut of the infant.^56,57^ Here we show that fucosylated HMOs support robust growth across all strains of *Akkermansia*. Growth did, however, vary by strain suggesting potential differences in growth and metabolic efficiencies across strains. When grown on 2’-FL, the liberated fucose was rapidly depleted from the culture medium, while the lactose component accumulated in the culture medium (except for CSUN-19) suggesting a general preference for fucose over lactose in *Akkermansia*. Cleavage of fucose from HMOs (and mucin) is mediated by fucosidases belonging to the GH29 or GH95 families,^31,38,49^ which were both found in all the four *Akkermansia* strains. GH141, a putative fucosidase or xylanase, was also observed in some of our AmI and AmIII genomes in this study. Kostopoulos et al. recently demonstrated that a GH29 gene product (encoded by Amuc_0010) in *A. muciniphila* Muc^T^ had relatively poor catalytic activity against 2’-FL, suggesting that 2’-FL was not the preferred substrate for this enzyme ^26^. Overall, *A. muciniphila* Muc^T^ has four GH29 gene annotations, two of GH95, and one of GH141, and all of these GH families could potentially encode enzymes that are involved in degradation of fucosylated HMOs containing the α1-2 linkage. The number of these same GH families also varied across phylogroups potentially leading the differences in growth efficiencies we observed. Given the prominent role of fucosylated HMOs in modulating the microbiome and enhancing health, and that the concentration of 2’-FL along with lacto-N-fucopentaose were highest during early lactation,^58^ the diversity of fucosidases available in each strain make *Akkermansia* a potential candidate to further investigate in the field of infant-associated probiotics.

Sialyl oligosaccharides are associated with many benefits to neonates and infants. ^59,60^ For example, Charbonneau and colleagues demonstrated that the concentration of sialylated HMOs in breastmilk correlated with growth in healthy Malawian infants ^60^. Furthermore, gnotobiotic mammals receiving fecal microbiota from infants with stunted growth and supplementation with sialylated bovine milk oligosaccharides, improved growth (measured as weight gain and bone mass), with their gut microbiota developing metabolic fitness evidenced by an increase in genes related to energy metabolism ^60^. Sialic acid is an essential component of brain gangliosides, and plays an important role in neuronal development, memory formation, and cognition.^59^ Three weeks of dietary supplementation with 3’-SL or 6’-SL administered to day-old piglets, increased the ganglioside-bound sialic acid in the brain of the piglets, thus providing essential nutrients for brain growth and neurodevelopment^61^. With regards to *Akkermansia* and sialylated HMOs, all four *Akkermansia* strains showed enhanced growth on 6’-SL and were able to deconstruct this sialylated oligosaccharide, but the growth yield and the percent of the substrate degraded varied significantly across strains. These differences in yield and degradation did not align with the sialidase (GH33) gene copy number. For example, strain CSUN-56 representing AmIII has 5 sialidase annotations, and exhibited relatively poor growth with little degradation of 6’-SL. This incongruence between the bacterial gene number of a GH metabolizing a substrate and the physiological response to that substrate indicates the need to examine the transcription of the GHs and the enzyme kinetics of associated GHs involved in the complete deconstruction of a substrate and their transport into the cell. However, the accumulation of sialic acid in spent medium after growth on 6’-SL in all strains agrees with previous reports of *Akkermansia* lacking the NAN operon for import and consumption of sialic acid.^26^ The sialic acid released from the non-reducing end of the sugars enables access to the remaining oligosaccharides, while also potentially encouraging the outgrowth of sialic acid metabolizing, abundantly-present commensal species such as *B. fragilis*, *Faecalibacterium prausnitzii*, *Ruminococcus gnavus*, and members of the *Lactobacillus* and *Bifidobacterium* genus.^17,62,63^ Several species of *Enterobacteriaceae* such as *Escherichia coli* and *Salmonella enterica* also thrive in a sialic acid-rich gut environment, with their fitness and virulence directly proportional to their ability to metabolize sialic acid.^62^ Interestingly though, studies in piglets demonstrated that supplementation with 6’-SL enhanced colonic bacteria such as *Collinsella aerofaciens Ruminococcus*, *Faecalibacterium*, and *Prevotella* spp., while suppressing *Enterobacteriaceae*, *Enterococcaceae*, *Lachnospiraceae*, and Lactobacillales.^61^ Given the vulnerability of the infant population and the immaturity of the gut microbiome in early life, identifying the metabolic fate of the sialic acid and the interaction between *Akkermansia* and sialic acid-metabolizing commensals and potential pathogens warrants further investigation.

*Akkermansia* are adapted to robust growth on mucin due to their habitation in the gut epithelial mucosa.^64^ Furthermore HMOs, that are resistant to host digestive enzymes, are presented to the colonic microbiota in a mucin-rich background of the infant gut.^65^ We therefore included mucin in our HMO utilization experiments. However, it is recognized that *Akkermansia* can grow in a mucin-deficient medium supplemented with GlcNAc, threonine, and tryptone.^9^ GlcNAc is a requirement for growth as *Akkermansia* do not express the enzyme required for conversion of fructose-6-phosphate to glucosamine-6-phosphate, an essential component of the cell wall peptidoglycan.^64^ GlcNac was thus added into the basal growth medium by Kostopoulos and colleagues. whilst investigating HMO utilization by *Akkermansia muciniphila* Muc^T^.^26^ We speculate that *Akkermansia* may potentially grow in the presence of GlcNAc-containing HMOs such as LNT or LNnT, provided that the amino acid sources are added to the growth medium. However, since our current technique precluded analysis of GlcNAc, further growth experiments and chemical analyses are required to confirm this prediction.

In conclusion, human-associated *Akkermansia* can utilize a variety of host-derived HMOs for growth *in vitro* in a strain-dependent manner. This implies that the prebiotic effects of HMOs will depend on the resident strain of *Akkermansia* present in an individual. When grown on HMO, *Akkermansia* liberate sugars and produce fermentation products that can fuel other members of the gut microbiome. Considering the presence of *Akkermansia* in the neonatal gut and the high abundance of oligosaccharides in mothers’ milk, *Akkermansia* may be considered as keystone species and nature’s way of engineering early life gut microbiome to grant long-lasting effects on metabolic fitness.

## Methods

### Recruitment and Sampling

Fecal samples used for *Akkermansia* isolations were obtained from 17 consenting healthy adults as previously described by Kirmiz et al.^10^ under protocol #1516-146, with approval from the Institutional Review Board at California State University, Northridge. Samples were refrigerated (4°C) and inoculated into culture medium (see below) within 24 h of collection.

### Bacterial isolation and identification

*Akkermansia* isolation and identification were conducted as previously described.^10^ Briefly, 5 mL of anaerobic Basal Mucin Medium (BMM) containing 0.5% v/v mucin (Supplementary Table S1) was inoculated with fecal swabs in serum tubes and a ten-fold serial dilution up to 10^-6^ or 10^-7^ was performed for each sample. Cultures were incubated at 37°C for up to 5 d, and those with oval cells in pairs were further diluted in broth medium and/or transferred to BMM agar until purity could be verified microscopically using a Zeiss Axioskop or as single colonies on BMM agar. For identification, genomic DNA was extracted using the DNAeasy^®^ UltraClean^®^ Microbial kit Isolation Kit (Qiagen Inc., MD, USA) and the near full-length 16S rRNA gene was amplified using primers 8F (5’-AGAGTTTGATCCTGGCTCAG-3’) and 1492R (5’-TACGGTTACCTTGTTACGA-3’) with the GoTaq^®^ Hot Start Colorless Master Mix (Promega Corp., Madison, WI, USA). PCR was performed using Eppendorf Vapo Protect Mastercycler Pro S 6325 (Hamburg, Germany) and included an activation/denaturation step at 95°C for 3 min, followed by 30 cycles of 95°C for 45 s, 45°C for 1 min and 72°C for 1 min 45s and a final extension step at 72°C for 7 min, followed by a hold at 4°C. PCR products were purified (QIAquick PCR Purification Kit, Qiagen Inc.) and sequenced using either the 8F or 1492R primer on an ABI Prism 3730 DNA sequencer (Laragen Sequencing and Genotyping, Culver City, CA). If sequences were pure and positively BLASTed to *A. muciniphila* in GenBank, the near full-length 16S rRNA gene was sequenced with additional primers (515F (GTGCCAGCMGCCGCGGTAA), 806R (GGACTACHVGGGTWTCTAAT), and 8F or 1492R). Sequences associated with each isolate were then assembled in Geneious 7.1.3 (https://www.geneious.com) and imported into ARB ^66^. In ARB, sequences were manually aligned with secondary structure constraints against the 16S rRNA gene sequence of *A. muciniphila* Muc^T^. To determine phylogroup affiliation based on 16S rRNA gene sequences, each isolate was added to our in-house database of *Akkermansia* 16S rRNA gene sequences as previously described.^10,15^ Masked alignments were exported from ARB and imported into Kumar and colleagues’^67^ MEGA7 where phylogenetic reconstruction was performed using the maximum-likelihood approach.

### Genome sequencing, assembly, annotation, and phylogenomics

Eleven *Akkermansia* isolates were selected for genome sequencing across three different sequencing efforts. DNA from strains CSUN-7 and CSUN-12 were sequenced according to the protocol described by Oliver and colleagues^68^ under the ‘Illumina sequencing’ section. To obtain enough DNA for this sequencing protocol, four 5 mL overnight BMM-grown cultures were extracted as described above and extracts were pooled and concentrated using ethanol precipitation with 3M sodium acetate. Illumina sequencing libraries were then prepared as described by Oliver and colleagues. DNA from strains CSUN-17, CSUN-19, CSUN-33, and CSUN-34 were sequenced according to the methods described by Parker and colleagues.^69^ For both this and the following sequencing efforts, enough quality genomic DNA was obtained from a single 5 mL BMM-grown culture of each isolate extracted as described above. The DNA from the remaining isolates (CSUN-37, CSUN-50, CSUN-56, CSUN-58, and CSUN-59) were sequenced on an Illumina NextSeq 550 (2×150bp) by the Microbial Genome Sequencing Center (Pittsburgh, PA, USA).

For assembly and annotation, paired fastq files of each isolate were submitted to PATRIC (v 3.6.3)^70^ for their “Comprehensive Genome Analysis” workflow that uses Unicycler^71^ to assemble genomes and RASTtk^72^ for annotation. For comparison, the nucleotide sequence file for *A. muciniphila* Muc^T^ ATCC BAA-835 (SAMN00138213) was downloaded from GenBank and annotated identical to the novel isolate genomes also using tools in PATRIC. To investigate the carbohydrate degrading potential of each *Akkermansia* phylogroup, the assembled contigs of the new isolates (n=11) were combined with 74 publicly available *Akkermansia* genomes ^10,15,73^ and submitted to the online dbCAN meta server for CAZyme annotation ^27–29^. dbCAN uses three tools - HMMER ^74^, DIAMOND ^75^, and Hotpep ^76^ - for automated CAZyme annotation. Annotations were considered only if they matched in at least two of the three tools. Individual count files were tabulated and compiled using a custom python script to generate a frequency table for all genomes (n=85). The resulting table was sorted and trimmed to include only glycoside hydrolase (GH) annotations and a heatmap was constructed in R ^77^ using the heatmap.2 function in the gplots library ^78^. Cluster dendrograms in the heatmap were calculated using average linkage hierarchical clustering based on Bray-Curtis dissimilarity matrices calculated using the vegan package also in R ^79^. To determine if there were differences in the number of GH predictions between phylogroups, a Kruskal-Wallis test (kruskal.test) followed by the Dunn’s test (dunn.test, method=‘bonferroni’) were performed in R.

For phylogenomic analysis, amino acid sequences of 49 ribosomal protein coding genes ^80^ were extracted and concatenated from assembled genomes using the ‘phylogenomics’ workflow in anvi’o ^81^. The concatenated fasta file was then imported into MEGA7 ^67^, aligned using MUSCLE ^82^, and a phylogenetic tree was made using the maximum-likelihood method ^83^ with 100 bootstraps.

### HMO growth experiments

To determine if *Akkermansia* strains could grow using HMOs, we performed a series of growth experiments in a customized medium, prepared by increasing the concentrations of threonine and tryptone (TT) in BMM ^9^. This medium, hereafter referred to as BMM-TT (supplementary Table S1), was supplemented with individual HMOs before inoculation with the chosen *Akkermansia* strains. Five HMOs were tested, namely 2’-FL, 3-FL, LNT, LNnT, and 6’-SL (Glycom, Hørsholm, Denmark). Lactose was also included in these growth experiments since it is the backbone of HMOs. Initially, representative isolates of each phylogroup (AmI = *A. muciniphila* Muc^T^, AmII = *Akkermansia* CSUN-17, AmIII = *Akkermansia* CSUN-56, and AmIV = *Akkermansia* CSUN-19; Supplementary Table S2) were grown overnight (18 to 24 h) in BMM at 37 °C under an atmosphere of N_2_/CO_2_ (70:30, vol/vol). Cultures were then standardized to an OD_600nm_ of 0.5 in fresh BMM and used to inoculate (10%) 200 μL of BMM-TT or BMM-TT supplemented with 20 mM of each HMO (or lactose) in 96-well microtiter plates (Falcon^®^, Corning Incorporated, NY, USA) in triplicate. Wells were overlaid with 30 μL of filter-sterilized mineral oil to prevent evaporation over the 48-h incubation period. After 48 h of anaerobic (N_2_/CO_2_/H_2_; 80:15:5 [vol/vol]) incubation at 37 °C in a Bactron IV anaerobic chamber (Sheldon Manufacturing, Inc., Cornelius, OR), plates were shaken for 10 s and OD_600nm_ was determined using a Spectramax microplate reader (Molecular Devices, San Jose, CA, USA). Growth was determined as ΔOD_600nm_ i.e., the change in OD_600nm_ growth in the BMM-TT supplemented with the HMOs relative to the growth in HMO-unsupplemented BMM-TT (i.e., BMM-TT + HMO OD_600nm_ – BMM-TT OD_600nm_). If OD_600nm_ were over 1.0, samples were diluted in half with a fresh medium and reread. Each experiment was conducted in triplicate and repeated at least two times. To test for differences in growth across strains, we used a repeated measures analysis of variance (ANOVA) followed by Tukey’s honestly significant differences (HSD) test as appropriate. Uninoculated controls were included in each experiment and remained negative for growth.

To verify the degradation of three HMOs (2’-FL, LNT, and 6’-SL), the above experiments were repeated in 1.5 mL of BMM-TT supplemented with 4 mM HMO. These experiments were conducted in 24-well microtiter plates (Costar, Corning Incorporated, NY, USA) sealed with Microseal^®^ ‘A’ Film (Bio-Rad Laboratories, Inc., Hercules, CA, USA) instead of mineral oil. Plates were incubated and OD_600nm_ and ΔOD_600nm_ were measured after 48 h as described above. For glycoanalytics, 0.5ml aliquots were taken at time 0 and 48 h after incubation, transferred to Eppendorf tubes and centrifuged at 10,000 × g for 3 min at 4°C. The cell-free supernatants were stored at −20°C for glycoanalytics as described below. To compare growth, statistical analysis was conducted as described above.

### HMO quantification

Culture supernatants were collected at time 0 and after 48 h of incubation to measure the degradation of 2’-FL, LNT, and 6’-SL. In addition to each parent HMO, individual sugars (with the exception of GlcNAc from LNT) of the three HMOs were also quantitatively measured using high-performance anion exchange chromatography with pulsed amperometric detection (HPAEC-PAD) ^84,85^. Frozen, cell-free spent culture media were thawed in a water bath, vortexed thoroughly to make a uniform mixture, and centrifuged at 7,000 × g for 5 min at 10°C and 1μL of the spent culture media was injected in HPAEC-PAD for detection of the above-mentioned sugars.

Carbohydrate analysis was done on Dionex-ICS3000 (Thermo Scientific, Sunnyvale, CA, USA) using CarboPac PA-1 column (4 mm x 250 mm) attached with Carbo PA1-guard column (4 mm x 50 mm). Detection of monosaccharides and oligosaccharides was done using standard Quad potential for carbohydrate analysis as supplied by the manufacturer. A gradient mixture of two solvents along with HPLC grade water was used for optimum separation of monosaccharides and oligosaccharides present in the sample. Solvent-A (Water), Solvent-B (100 mM NaOH + 7 mM NaOAc) and Solvent-C (100 mM NaOH + 250 mM NaOAc) were used as elution solvents at a flow rate of 1.0 mL/min. Gradient mixture details are given in Supplementary Table S4. Sugars were quantified by comparing with the area under the peaks from a standard mixture of fucose, galactose, glucose, 3-FL, lactose, 2’-FL, LNnT, LNT, sialic acid (Neu5Ac), 6’-SL, and 3-SL. Representative chromatograms are presented in Supplemental Figure S1. To determine the percent of HMOs utilized, the amount remaining after 48 h of incubation was divided by the amount at time 0 and multiplied by [100-(HMO 48 h/HMO 0 h)] *100.

## Acknowledgements

The authors would like to thank Louise Vigsnaes and Glycom for their generous donation of HMOs and their continued support of our work.

## Author Contributions

E.L. helped conceive the project, conducted all HMO growth experiments, aided in data interpretation, and helped write the manuscript. S.G.P. performed statistical analysis, aided in data interpretation, and helped write the manuscript. N.K. helped conceive the project and write the manuscript. S.H., E.H., M.H., A.K., C.M., K.O., L.P., K.R., and P.S. helped collect samples and isolate *Akkermansia* strains. E.S. helped with the genomic analysis. B.C. and M.P. performed the glycoanalytics and helped with data interpretation. C.T.P and K.C. performed genome sequencing and helped with genomic analysis. G.E.F. conceived of the project, aided in data interpretation, performed genomic and statistical analysis, and helped write the paper.

## Supplementary material

Supplemental data for this article can be accessed on the publisher’s website.

## Data availability statement

The data that support the findings of this study are openly available in NCBI BioProject database at https://www.ncbi.nlm.nih.gov/bioproject/609771, accession number [PRJNA609771].

## References

1. Derrien M, Vaughan EE, Plugge CM, de Vos WM. *Akkermansia muciniphila* gen. nov., sp. nov., a human intestinal mucin-degrading bacterium. Int J Syst Evol Microbiol. 2004; 54:1469–76.

2. Kim S, Shin Y-C, Kim T-Y, Kim Y, Lee Y-S, Lee S-H, et al. Mucin degrader *Akkermansia muciniphila* accelerates intestinal stem cell-mediated epithelial development. Gut Microbes 2021; 13:1–20.

3. Dao MC, Everard A, Aron-Wisnewsky J, Sokolovska N, Prifti E, Verger EO, et al. *Akkermansia muciniphila* and improved metabolic health during a dietary intervention in obesity: relationship with gut microbiome richness and ecology. Gut 2016; 65:426–36.

4. Rajilic-Stojanovic M, Shanahan F, Guarner F, de Vos WM. Phylogenetic analysis of dysbiosis in ulcerative colitis during remission. Inflamm Bowel Dis. 2013; 19:481–8.

5. Png CW, Linden SK, Gilshenan KS, Zoetendal EG, McSweeney CS, Sly LI, et al. Mucolytic bacteria with increased prevalence in IBD mucosa augment *in vitro* utilization of mucin by other bacteria. Am J Gastroenterol. 2010; 105:2420–8.

6. Lee M-J, Kang M-J, Lee S-Y, Lee E, Kim K, Won S, et al. Perturbations of gut microbiome genes in infants with atopic dermatitis according to feeding type. J Allergy Clin Immunol. 2018; 141:1310–9.

7. Ouwerkerk JP, van der Ark KCH, Davids M, Claassens NJ, Finestra TR, de Vos WM, et al. Adaptation of *Akkermansia muciniphila* to the oxic-anoxic interface of the mucus layer. Appl Environ Microbiol. 2016; 82:6983–93.

8. Robbe C, Capon C, Coddeville B, Michalski JC. Structural diversity and specific distribution of O-glycans in normal human mucins along the intestinal tract. Biochem J. 2004; 384:307–16.

9. Ottman N, Davids M, Suarez-Diez M, Boeren S, Schaap PJ, Martins Dos Santos VAP, et al. Genome-scale model and omics analysis of metabolic capacities of *Akkermansia muciniphila* reveal a preferential mucin-degrading lifestyle. Appl Environ Microbiol. 2017; 83.

10. Kirmiz N, Galindo K, Cross KL, Luna E, Rhoades N, Podar M, et al. Comparative genomics guides elucidation of vitamin B12 biosynthesis in novel human-associated *Akkermansia* strains. Appl Environ Microbiol. 2020; 86:e02117–19.

11. Chia LW, Hornung BVH, Aalvink S, Schaap PJ, de Vos WM, Knol J, et al. Deciphering the trophic interaction between *Akkermansia muciniphila* and the butyrogenic gut commensal *Anaerostipes caccae* using a metatranscriptomic approach. Antonie Van Leeuwenhoek 2018; 111:859–73.

12. Cani PD, Van Hul M, Lefort C, Depommier C, Rastelli M, Everard A. Microbial regulation of organismal energy homeostasis. Nat Metab. 2019; 1:34–46.

13. Ottman N, Reunanen J, Meijerink M, Pietila TE, Kainulainen V, Klievink J, et al. Pililike proteins of *Akkermansia muciniphila* modulate host immune responses and gut barrier function. PLoS ONE 2017; 12:e0173004.

14. Plovier H, Everard A, Druart C, Depommier C, Van Hul M, Geurts L, et al. A purified membrane protein from A*kkermansia muciniphila* or the pasteurized bacterium improves metabolism in obese and diabetic mice. Nat Med. 2017; 23:107–13.

15. Guo X, Li S, Zhang J, Wu F, Li X, Wu D, et al. Genome sequencing of 39 *Akkermansia muciniphila* isolates reveals its population structure, genomic and functional diverisity, and global distribution in mammalian gut microbiotas. BMC Genom. 2017; 18:800.

16. Bode L. Human milk oligosaccharides: every baby needs a sugar mama. Glycobiology 2012; 22:1147–62.

17. Coker JK, Moyne O, Rodionov DA, Zengler K. Carbohydrates great and small, from dietary fiber to sialic acids: How glycans influence the gut microbiome and affect human health. Gut Microbes 2021; 13:1869502.

18. Bode L, Jantscher-Krenn E. Structure-function relationships of human milk oligosaccharides. Adv Nutr. 2012; 3:383s–91s.

19. Kunz C, Rudloff S, Baier W, Klein N, Strobel S. Oligosaccharides in human milk: structural, functional, and metabolic aspects. Annu Rev Nutr. 2000; 20:699–722.

20. Garrido D, Barile D, Mills DA. A molecular basis for bifidobacterial enrichment in the infant gastrointestinal tract. Adv Nutr. 2012; 3:415S–21S.

21. Kuntz S, Kunz C, Rudloff S. Oligosaccharides from human milk induce growth arrest via G2/M by influencing growth-related cell cycle genes in intestinal epithelial cells. Br J Nutr. 2009; 101:1306–15.

22. Morrow AL, Ruiz-Palacios GM, Jiang X, Newburg DS. Human-milk glycans that inhibit pathogen binding protect breast-feeding infants against infectious diarrhea. J Nutr. 2005; 135:1304–7.

23. Ward RE, Niñonuevo M, Mills DA, Lebrilla CB, German JB. *In vitro* fermentation of breast milk oligosaccharides by *Bifidobacterium infantis* and *Lactobacillus gasseri*. Appl Environ Microbiol. 2006; 72:4497–9.

24. Marcobal A, Barboza M, Froehlich JW, Block DE, German JB, Lebrilla CB, et al. Consumption of human milk oligosaccharides by gut-related microbes. J Agric Food Chem. 2010; 58:5334–40.

25. Yu ZT, Chen C, Newburg DS. Utilization of major fucosylated and sialylated human milk oligosaccharides by isolated human gut microbes. Glycobiology 2013; 23:1281–92.

26. Kostopoulos I, Elzinga J, Ottman N, Klievink JT, Blijenberg B, Aalvink S, et al. *Akkermansia muciniphila* uses human milk oligosaccharides to thrive in the early life conditions *in vitro*. Sci Rep. 2020; 10:14330.

27. Cantarel BL, Coutinho PM, Rancurel C, Bernard T, Lombard V, Henrissat B. The Carbohydrate-Active EnZymes database (CAZy): an expert resource for glycogenomics. Nucleic Acids Res. 2009; 37:D233–8.

28. Yin Y, Mao X, Yang J, Chen X, Mao F, Xu Y. dbCAN: a web resource for automated carbohydrate-active enzyme annotation. Nucleic Acids Res. 2012; 40:W445–W51.

29. Zhang H, Yohe T, Huang L, Entwistle S, Wu P, Yang Z, et al. dbCAN2: a meta server for automated carbohydrate-active enzyme annotation. Nucleic Acids Res. 2018; 46:W95–W101.

30. Tailford LE, Crost EH, Kavanaugh D, Juge N. Mucin glycan foraging in the human gut microbiome. Front Genet. 2015; 6.

31. Katoh T, Ojima MN, Sakanaka M, Ashida H, Gotoh A, Katayama T. Enzymatic adaptation of *Bifidobacterium bifidum* to host glycans, viewed from glycoside hydrolyases and carbohydrate-binding modules. Microorganisms 2020; 8.

32. Low KE, Smith SP, Abbott DW, Boraston AB. The glycoconjugate-degrading enzymes of *Clostridium perfringens*: Tailored catalysts for breaching the intestinal mucus barrier. Glycobiology 2020.

33. Ioannou A, Knol J, Belzer C. Microbial glycoside hydrolases in the first year of life: an analysis review on their presence and importance in infant gut. Front Microbiol. 2021; 12:1345.

34. Chen H, Kim J, Kendall DA. Competition between functional signal peptides demonstrates variation in affinity for the secretion pathway. J Bacteriol. 1996; 178:6658–64.

35. Owji H, Nezafat N, Negahdaripour M, Hajiebrahimi A, Ghasemi Y. A comprehensive review of signal peptides: Structure, roles, and applications. Eur J Cell Biol. 2018; 97:422–41.

36. Zhai Q, Feng S, Arjan N, Chen W. A next generation probiotic, *Akkermansia muciniphila*. Crit Rev Food Sci Nutr. 2019; 59:3227–36.

37. Becken B, Davey L, Middleton DR, Mueller KD, Sharma A, Holmes ZC, et al. Genotypic and phenotypic diversity among human isolates of *Akkermansia muciniphila*. mBio 2021; 12:e00478–21.

38. Sela DA, Garrido D, Lerno L, Wu S, Tan K, Eom H-J, et al. *Bifidobacterium longum* subsp. *infantis* ATCC 15697 α-fucosidases are active on fucosylated human milk oligosaccharides. Appl Environ Microbiol. 2012; 78:795–803.

39. Kiyohara M, Tanigawa K, Chaiwangsri T, Katayama T, Ashida H, Yamamoto K. An exo-alpha-sialidase from bifidobacteria involved in the degradation of sialyloligosaccharides in human milk and intestinal glycoconjugates. Glycobiology 2011; 21:437–47.

40. Sela DA, Li Y, Lerno L, Wu S, Marcobal AM, German JB, et al. An infant-associated bacterial commensal utilizes breast milk sialyloligosaccharides. J Biol Chem. 2011; 286:11909–18.

41. Matsuki T, Yahagi K, Mori H, Matsumoto H, Hara T, Tajima S, et al. A key genetic factor for fucosyllactose utilization affects infant gut microbiota development. Nat Commun. 2016; 7:11939.

42. James K, Motherway MO, Bottacini F, van Sinderen D. *Bifidobacterium breve* UCC2003 metabolises the human milk oligosaccharides lacto-N-tetraose and lacto-N-neo-tetraose through overlapping, yet distinct pathways. Sci Rep. 2016; 6:38560.

43. Garrido D, Ruiz-Moyano S, Kirmiz N, Davis JC, Totten SM, Lemay DG, et al. A novel gene cluster allows preferential utilization of fucosylated milk oligosaccharides in *Bifidobacterium longum* subsp. *longum* SC596. Sci Rep. 2016; 6:35045.

44. Marcobal A, Barboza M, Sonnenburg ED, Pudlo N, Martens EC, Desai P, et al. *Bacteroides* in the infant gut consume milk oligosaccharides via mucus-utilization pathways. Cell Host Microbe 2011; 10:507–14.

45. Garrido D, Kim JH, German JB, Raybould HE, Mills D. Oligosaccharide binding proteins from *Bifidobacterium longum* subsp. *infantis* reveal a preference for host glycans. PLoS ONE 2011; 6.

46. Yoshida E, Sakurama H, Kiyohara M, Nakajima M, Kitaoka M, Ashida H, et al. *Bifidobacterium longum* subsp. *infantis* uses two different β-galactosidases for selectively degrading type-1 and type-2 human milk oligosaccharides. Glycobiology 2012; 22:361–8.

47. Ruiz-Moyano S, Totten SM, Garrido D, Smilowitz JT, German JB, Lebrilla CB. Variation in consumption of human milk oligosaccharides by infant gut-associated strains of *Bifidobacterium breve*. Appl Environ Microbiol. 2013; 79.

48. Turroni F, Bottacini F, Foroni E, Mulder I, Kim JH, Zomer A, et al. Genome analysis of *Bifidobacterium bifidum* PRL2010 reveals metabolic pathways for host-derived glycan foraging. Proc Natl Acad Sci U S A. 2010; 107:19514–9.

49. Marcobal A, Sonnenburg JL. Human milk oligosaccharide consumption by intestinal microbiota. Clin Microbiol Infect. 2012; 18:12–5.

50. Sampson EM, Bobik TA. Microcompartments for B12-dependent 1,2-propanediol degradation provide protection from dna and cellular damage by a reactive metabolic intermediate. J Bacteriol. 2008; 190:2966–71.

51. Engels C, Ruscheweyh H-J, Beerenwinkel N, Lacroix C, Schwab C. The common gut microbe *Eubacterium hallii* also contributes to intestinal propionate formation. Front Microbiol. 2016; 7.

52. Yatsunenko T, Rey FE, Manary MJ, Trehan I, Dominguez-Bello MG, Contreras M, et al. Human gut microbiome viewed across age and geography. Nature 2012; 486:222–7.

53. Collado MC, Derrien M, Isolauri E, de Vos WM, Salminen S. Intestinal Integrity and *Akkermansia muciniphila*, a mucin-degrading member of the intestinal microbiota present in infants, adults, and the elderly. Appl Environ Microbiol. 2007; 73:7767–70.

54. Collado MC, Laitinen K, Salminen S, Isolauri E. Maternal weight and excessive weight gain during pregnancy modify the immunomodulatory potential of breast milk. Pediatr Res. 2012; 72:77–85.

55. Lackey KA, Williams JE, Meehan CL, Zachek JA, Benda ED, Price WJ, et al. What’s normal? Microbiomes in human milk and infant feces are related to each other but vary geographically: The INSPIRE study. Front Nutr. 2019; 6:45.

56. Korpela K, Salonen A, Hickman B, Kunz C, Sprenger N, Kukkonen K, et al. Fucosylated oligosaccharides in mother’s milk alleviate the effects of caesarean birth on infant gut microbiota. Sci Rep. 2018; 8:13757.

57. Aakko J, Kumar H, Rautava S, Wise A, Autran C, Bode L, et al. Human milk oligosaccharide categories define the microbiota composition in human colostrum. Benef Microbes 2017; 8:563–7.

58. Thurl S, Munzert M, Henker J, Boehm G, Müller-Werner B, Jelinek J, et al. Variation of human milk oligosaccharides in relation to milk groups and lactational periods. Br J Nutr 2010; 104:1261–71.

59. Wang B, Brand-Miller J. The role and potential of sialic acid in human nutrition. Eur J Clin Nutr. 2003; 57:1351–69.

60. Charbonneau MR, O’Donnell D, Blanton LV, Totten SM, Davis JCC, Barratt MJ, et al. Sialylated Milk Oligosaccharides Promote Microbiota-Dependent Growth in Models of Infant Undernutrition. Cell 2016; 164:859–71.

61. Jacobi SK, Yatsunenko T, Li D, Dasgupta S, Yu RK, Berg BM, et al. Dietary isomers of sialyllactose increase ganglioside sialic acid concentrations in the corpus callosum and cerebellum and modulate the colonic microbiota of formula-fed piglets. J Nutr. 2016; 146:200–8.

62. Almagro-Moreno S, Boyd EF. Bacterial catabolism of nonulosonic (sialic) acid and fitness in the gut. Gut Microbes 2010; 1:45–50.

63. Egan M, O’Connell Motherway M, Ventura M, van Sinderen D. Metabolism of sialic acid by *Bifidobacterium breve* UCC2003. Appl Environ Microbiol. 2014; 80:4414–26.

64. van der Ark KCH, Aalvink S, Suarez-Diez M, Schaap PJ, de Vos WM, Belzer C. Model-driven design of a minimal medium for *Akkermansia muciniphila* confirms mucus adaptation. Microb Biotechnol. 2018; 11:476–85.

65. Rokhsefat S, Lin A, Comelli EM. Mucin–Microbiota Interaction During Postnatal Maturation of the Intestinal Ecosystem: Clinical Implications. Dig Dis Sci. 2016; 61:1473–86.

66. Ludwig W, Strunk O, Westram R, Richter L, Meier H, Yadhukumar, et al. ARB: a software environment for sequence data. Nucleic Acids Res. 2004; 32:1363–71.

67. Kumar S, Stecher G, Tamura K. MEGA7: Molecular Evolutionary Genetics Analysis Version 7.0 for Bigger Datasets. Mol Biol Evol. 2016; 33:1870–4.

68. Oliver A, Kay M, Cooper KK. Comparative genomics of cocci-shaped *Sporosarcina* strains with diverse spatial isolation. BMC Genom. 2018; 19:310.

69. Parker CT, Cooper KK, Huynh S, Smith TP, Bono JL, Cooley M. Genome sequences of eight shiga toxin-producing *Escherichia coli* strains isolated from a produce-growing region in California. Microbiol Resour Announc. 2018; 7.

70. Wattam AR, Davis JJ, Assaf R, Boisvert S, Brettin T, Bun C, et al. Improvements to PATRIC, the all-bacterial bioinformatics database and analysis resource center. Nucleic Acids Res. 2017; 45:D535–d42.

71. Wick RR, Judd LM, Gorrie CL, Holt KE. Unicycler: Resolving bacterial genome assemblies from short and long sequencing reads. PLoS Comput Biol. 2017; 13:e1005595.

72. Brettin T, Davis JJ, Disz T, Edwards RA, Gerdes S, Olsen GJ, et al. RASTtk: a modular and extensible implementation of the RAST algorithm for building custom annotation pipelines and annotating batches of genomes. Sci Rep. 2015; 5:8365.

73. van Passel MW, Kant R, Zoetendal EG, Plugge CM, Derrien M, Malfatti SA, et al. The genome of *Akkermansia muciniphila*, a dedicated intestinal mucin degrader, and its use in exploring intestinal metagenomes. PLoS ONE 2011; 6:e16876.

74. Finn RD, Clements J, Eddy SR. HMMER web server: interactive sequence similarity searching. Nucleic Acids Res. 2011; 39:W29–37.

75. Buchfink B, Xie C, Huson DH. Fast and sensitive protein alignment using DIAMOND. Nat Methods 2015; 12:59–60.

76. Busk PK, Pilgaard B, Lezyk MJ, Meyer AS, Lange L. Homology to peptide pattern for annotation of carbohydrate-active enzymes and prediction of function. BMC Bioinform. 2017; 18:214.

77. R Core Team. R: A language and environment for statistical computing. Vienna, Austria: R Foundation for Statistical Computing, 2013.

78. Warnes G, Bolker B, Bonebakker L, Gentleman R, Huber W, Liaw A, et al. gplots: Various R programming tools for plotting data. https://cran.r-project.org/web/packages/gplots/index.html, 2009.

79. Oksanen J, Guillaume Blanchet F, Friendly M, Kindt R, Legendre P, McGlinn D, et al. vegan: community ecology package. In: 2.4-3. Rpv, ed. https://CRAN.R-project.org/package=vegan, 2017.

80. Campbell JH, O’Donoghue P, Campbell AG, Schwientek P, Sczyrba A, Woyke T, et al. UGA is an additional glycine codon in uncultured SR1 bacteria from the human microbiota. Proc Natl Acad Sci U S A. 2013; 110:5540–5.

81. Eren AM, Esen OC, Quince C, Vineis JH, Morrison HG, Sogin ML, et al. Anvi’o: an advanced analysis and visualization platform for ’omics data. PeerJ 2015; 3:e1319.

82. Edgar RC. MUSCLE: multiple sequence alignment with high accuracy and high throughput. Nucleic Acids Res. 2004; 32:1792–7.

83. Jones DT, Taylor WR, Thornton JM. The rapid generation of mutation data matrices from protein sequences. Comput Appl Biosci. 1992; 8:275–82.

84. Hardy MR, Townsend RR, Lee YC. Monosaccharide analysis of glycoconjugates by anion exchange chromatography with pulsed amperometric detection. Anal Biochem. 1988; 170:54–62.

85. Townsend RR, Hardy MR, Cumming DA, Carver JP, Bendiak B. Separation of branched sialylated oligosaccharides using high-pH anion-exchange chromatography with pulsed amperometric detection. Anal Biochem. 1989; 182:1–8.

